# AI-driven Deep Visual Proteomics defines cell identity and heterogeneity

**DOI:** 10.1101/2021.01.25.427969

**Authors:** Andreas Mund, Fabian Coscia, Réka Hollandi, Ferenc Kovács, András Kriston, Andreas-David Brunner, Michael Bzorek, Soraya Naimy, Lise Mette Rahbek Gjerdrum, Beatrice Dyring-Andersen, Jutta Bulkescher, Claudia Lukas, Christian Gnann, Emma Lundberg, Peter Horvath, Matthias Mann

**Author notes:** Authors contributed equally (Andreas Mund, Fabian Coscia).

## Abstract

The systems-wide analysis of biomolecules in time and space is key to our understanding of cellular function and heterogeneity in health and disease^1^. Remarkable technological progress in microscopy and multi-omics technologies enable increasingly data-rich descriptions of tissue heterogeneity^2,3,4,5^. Single cell sequencing, in particular, now routinely allows the mapping of cell types and states uncovering tremendous complexity^6^. Yet, an unaddressed challenge is the development of a method that would directly connect the visual dimension with the molecular phenotype and in particular with the unbiased characterization of proteomes, a close proxy for cellular function. Here we introduce Deep Visual Proteomics (DVP), which combines advances in artificial intelligence (AI)-driven image analysis of cellular phenotypes with automated single cell laser microdissection and ultra-high sensitivity mass spectrometry^7^. DVP links protein abundance to complex cellular or subcellular phenotypes while preserving spatial context. Individually excising nuclei from cell culture, we classified distinct cell states with proteomic profiles defined by known and novel proteins. AI also discovered rare cells with distinct morphology, whose potential function was revealed by proteomics. Applied to archival tissue of salivary gland carcinoma, our generic workflow characterized proteomic differences between normal-appearing and adjacent cancer cells, without admixture of background from unrelated cells or extracellular matrix. In melanoma, DVP revealed immune system and DNA replication related prognostic markers that appeared only in specific tumor regions. Thus, DVP provides unprecedented molecular insights into cell and disease biology while retaining spatial information.

## The deep visual proteomics concept

The versatility, resolution and multi-modal nature of modern microscopy delivers increasingly detailed images of single cell heterogeneity and tissue organization^8^. However, a pre-defined subset of proteins is usually targeted, far short of the actual complexity of the proteome. Taking advantage of recent developments in mass spectrometry (MS)-based technology, especially dramatically increased sensitivity, we here set out to enable the analysis of proteomes within their native, subcellular context to explore their contribution to health and disease. We developed a concept called Deep Visual Proteomics (DVP) that combines high-resolution imaging, artificial intelligence (AI)-based image analysis for single-cell phenotyping and isolation with a novel ultra-sensitive proteomics workflow^7^ (Fig. 1). A key challenge in realizing the DVP concept turned out to be the accurate definition of single cell boundaries and cell classes as well as the transfer of the AI defined features into proteomic samples, ready for analysis. To this end, we introduce the software ‘BIAS’ (Biology Image Analysis Software), which coordinates scanning and laser microdissection microscopes. This seamlessly combines data-rich imaging of cell cultures or archived biobank tissues (formalin-fixed and paraffin-embedded, FFPE) with deep learning-based cell segmentation and machine learning-based identification of cell types and states. Cellular or subcellular objects of interest are selected by the AI alone or after instruction and subjected to automated laser microdissection and proteomic profiling. Data generated by DVP can be mined to discover novel protein signatures providing molecular insights into proteome variation at the phenotypic level with full spatial meta-information. We show below that this concept provides a powerful, multi-layered resource for researchers with applications ranging from functional characterization of single cell heterogeneity to spatial proteomic characterization of disease tissues with the aim of assisting clinical decision-making.

**Fig.1:**
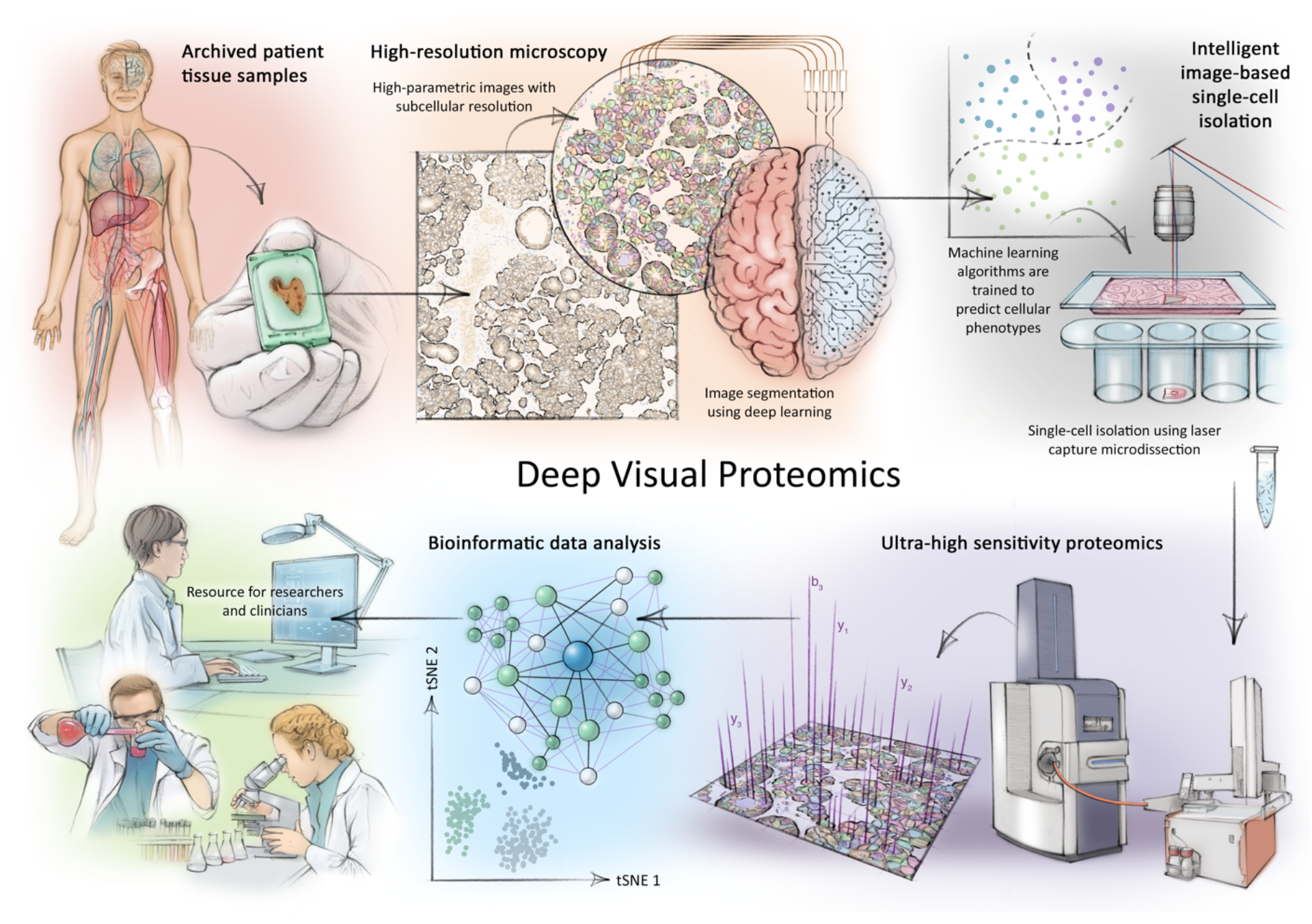
Deep Visual Proteomics concept and workflow. Deep Visual Proteomics (DVP) combines high-resolution imaging, artificial intelligence (AI)-guided image analysis for single-cell classification and isolation with a novel ultra-sensitive proteomics workflow7. DVP links data-rich imaging of cell culture or archived patient biobank tissues with deep learning-based cell segmentation and machine learning based identification of cell types and states. (Un)supervised AI-classified cellular or subcellular objects of interests undergo automated laser microdissection and mass spectrometry (MS)-based proteomic profiling. Subsequent bioinformatic data analysis enables data mining to discover protein signatures providing molecular insights into proteome variation in health and disease states at the level of single cells. DVP serves as resource for researchers and clinicians.

## The image processing and single cell isolation workflow

The microscopy-related aspects of the DVP workflow build on state-of-the-art high-resolution and whole-slide imaging as well as machine learning and deep learning (ML and DL) for image analysis. For the required pipeline, further advances in our image analysis software were needed, as well as downstream, automated, rapid single-cell laser microdissection.

First, we used scanning microscopy to obtain high-resolution whole-slide images and developed a software suite for integrative image analysis termed ‘BIAS’ (Methods). BIAS allows the processing of multiple 2D and 3D microscopy image file formats, supporting the major microscope vendors and data formats. It combines image preprocessing, deep learning-based image segmentation, feature extraction and machine learning-based phenotype classification. Building on a novel deep learning-based algorithm for cytoplasm and nucleus segmentation^9^, we undertook several optimizations to implement pre-processing algorithms to maintain high quality images across large image datasets. Deep learning methods require large training datasets, a major challenge due to the limited size of high-quality training data^10^. To address this challenge, we used NucleAIzer^9^ and applied project-specific image style transfer to synthetize artificial microscopy images resembling real images. This approach is inherently adaptable to different biological scenarios such as new cell and tissue types or staining techniques. We trained a deep learning model with these synthetic images for specific segmentation of the cellular compartment of interest (e.g. nucleus or cytoplasm, Fig. 2A), and benchmarked it against two leading deep learning approaches: unet4nuclei^11^ and Cellpose^12^ and a widely-used adaptive threshold- and object splitting-based method^13^. Notably, our deep learning algorithms for cell and nucleus segmentation of cell cultures and tissues showed the highest accuracy (Fig. 2A, Suppl. S1). For interactive cellular phenotype discovery, BIAS performs phenotypic feature extraction taking into account morphology and neighborhood features based on supervised and unsupervised machine-learning (Fig 2B, Methods). Importantly, we can combine feature-based phenotypic classification with biomarker expression levels from antibody staining for precise cell classification.

**Figure 2.**
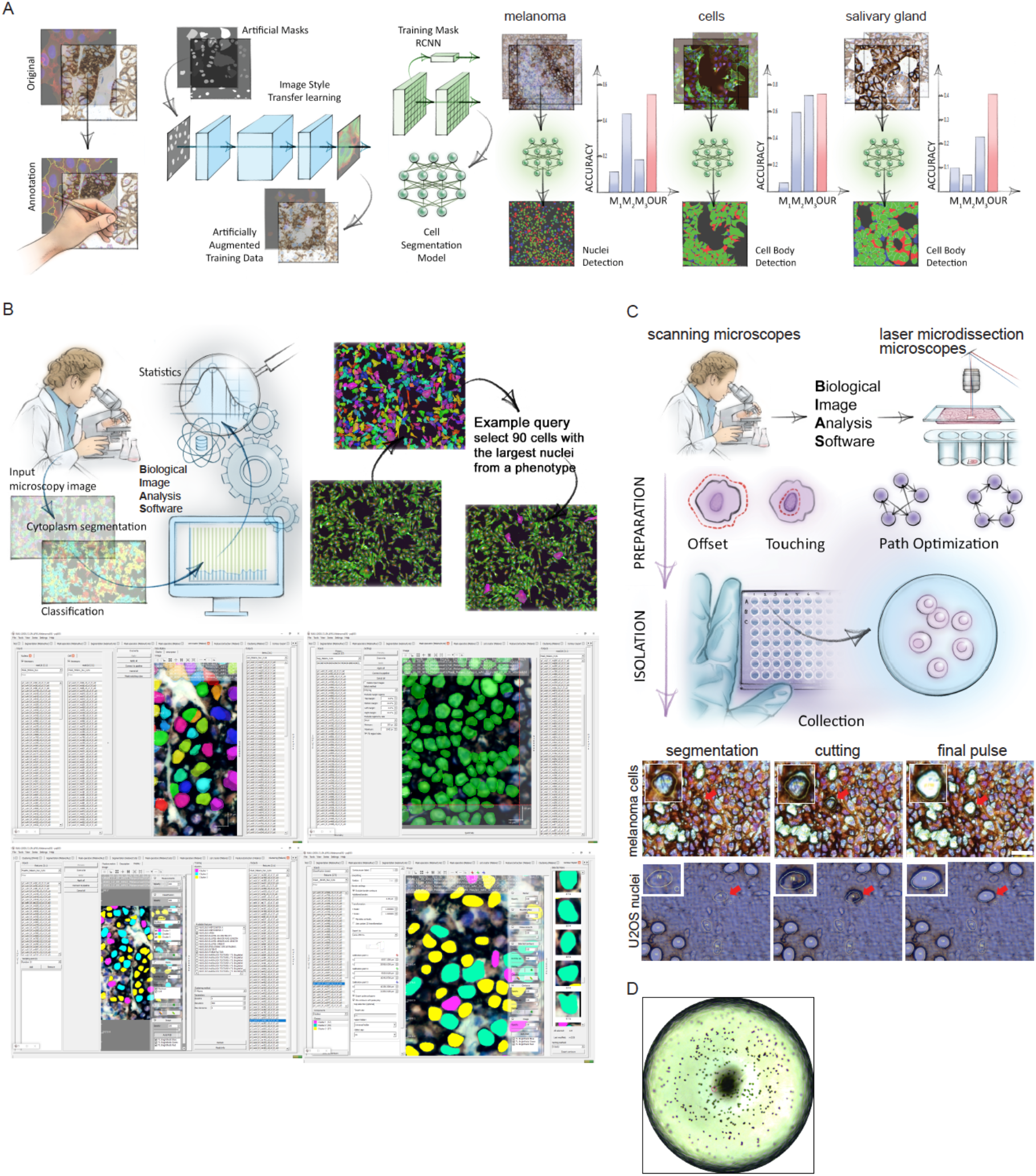
BIAS for integrative image analysis and automated LMD single-cell isolation. **A.** Left: AI-driven nuclei and cytoplasm segmentation of normal appearing and cancer cells and tissue using image style transfer learning in the Biological Image Analysis Software (BIAS), developed here. Right: We benchmarked the accuracy of its segmentation approach using the F1 metric and compared results to three additional methods M1-M3. Visual representation of the segmentation results: green areas correspond to true positive, blue to false positive and red to false negative. **B.** BIAS allows the processing of multiple 2D and 3D microscopy image file formats. Examples for image preprocessing, deep learning-based image segmentation, feature extraction and machine learning-based phenotype classification. **C.** BIAS also serves as the interface between the scanning and a laser microdissection microscope, allowing high accuracy transfers of cell contours between the microscopes. Upper panel: conceptual overview of cutting functions, cutting offset with respect to the object of interest and optimal path finding. Lower panel: Practical illustration of the functions in the upper panel. **D.** Captured single nuclei can be quality controlled in collection wells

To physically extract the cellular features discovered with BIAS, we developed an interface between scanning and laser microdissection microscopes (currently ZEISS PALM MicroBeam and Leica LMD6 & 7) (Fig. 2C). BIAS transfers cell contours between the microscopes, preserving full accuracy. Laser microdissection has a theoretical accuracy of 70 nm using a 150x objective and in practice we reached 200 nm. After optimization the LMD7 allows the excision of 700 collected high-resolution contours per hour, with full remote and automated operation (Methods). To prevent potential laser-induced damage, contours can be excised with a definable offset (Fig. 2C, D, video 1,2). In summary, BIAS successfully unifies scanning and laser microdissection microscopy on the basis of AI-driven image analysis.

## DVP defines single cell heterogeneity at the subcellular level

To determine if DVP can characterize functional differences between phenotypically distinct cells, we applied our workflow to an unperturbed cancer cell line (FUCCI - U2OS cells^14^). After deep learning-based segmentation for nuclei and cell membrane detection, we isolated 80-100 single cells or 250-300 nuclei per experiment (Fig. 2A, 3A, B). Although the analysis of small numbers of tissue cells by MS has been a long-standing goal, transfer, processing and analysis of these minute samples pose formidable analytical challenges^15^ which we addressed in turn. We processed samples on the basis of a recently developed workflow for ultra-low sample input^7,16^, that omits any sample transfer steps and ensures decrosslinking in very low volumes (Methods). We found that samples could be analyzed directly from 384 wells without any additional sample transfer or clean-up. MS measurements were performed with a data independent acquisition method using the parallel accumulation – serial fragmentation acquisition method an additional ion mobility dimension and optimal fragment (diaPASEF) ion usage on a newly developed mass spectrometer^17,7^. Replicates of cell and nucleus proteomes demonstrated a robust workflow with high quantitative reproducibility (Pearson r = 0.96). Proteomes of whole cells were very different from those of nuclei alone, as in subcellular proteomics experiments based on biochemical separation^18^ (Fig. 3C, Fig. S2A). This was likewise reflected in the bioinformatic enrichment analysis, with terms like plasma membrane, mitochondrion, nucleosomes and transcription factor complexes being highly significant (FDR < 10^−5^) (Fig. 3D).

**Fig.3:**
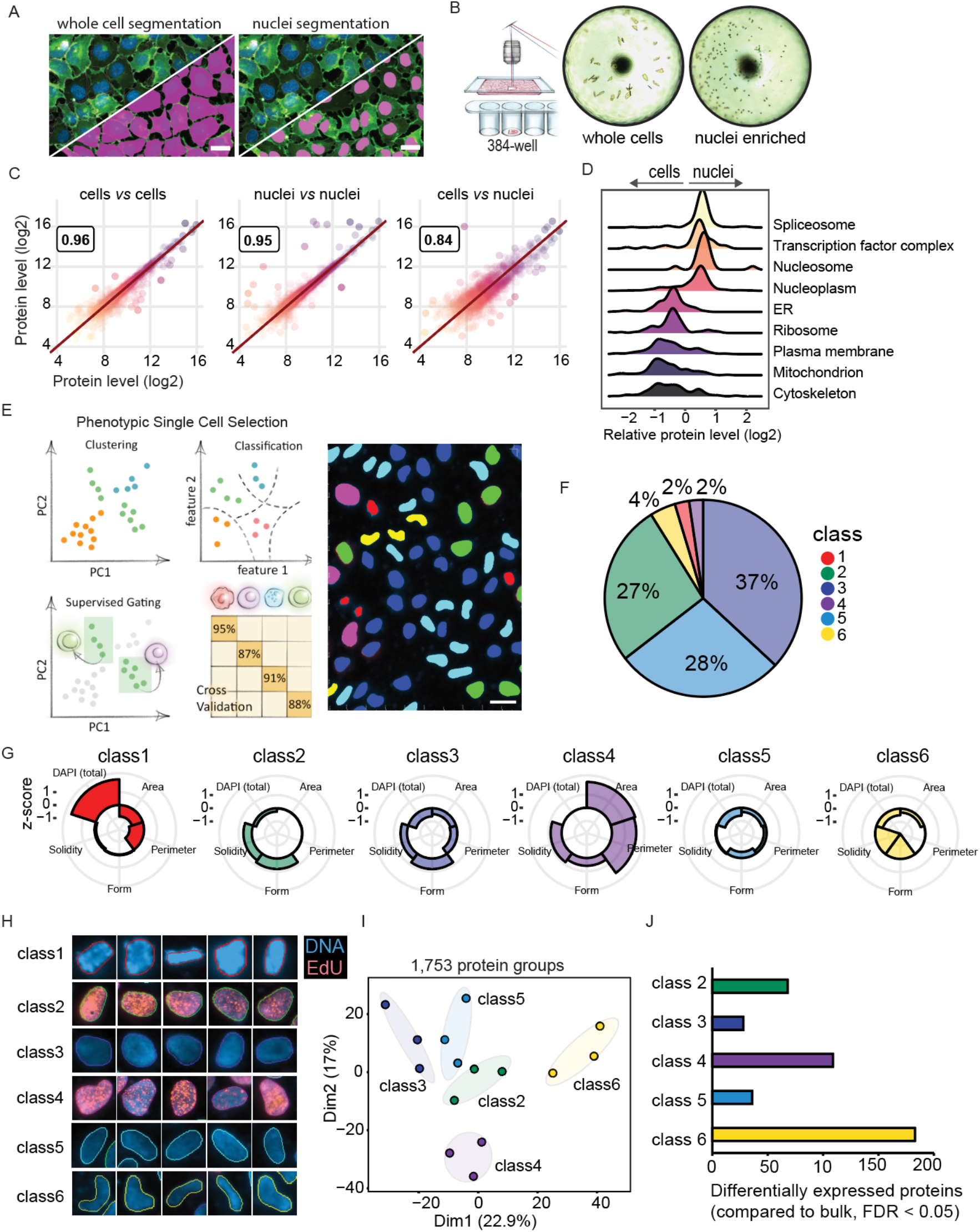
DVP defines single cell heterogeneity at the subcellular level. **A.** Deep learning-based segmentation of whole cells and nuclei in BIAS of DNA (DAPI) stained U2OS FUCCI cells. Scale bar =20μm **B.** Automated laser microdissection of whole cells and nuclei into 384-well plates. Images show wells after collection. **C.** Quantitative proteomic results of whole cell and nuclei replicates, and comparison between whole cells and nuclei. **D.** Relative protein levels (X-axis) of major cellular compartments between whole cell and nuclei specific proteomes. Y-axis displays point density. **E.** Left: Conceptual workflows of the phenotype finder model of BIAS for machine learning-based classification of cellular phenotypes. Right: Results of unsupervised ML-based classification of six distinct U2OS nuclei classes based on morphological features and DNA staining intensity. Colors represent classes. Scale bar = 20μm. **F.** Relative proportions of the six nuclei classes. **G.** Phenotypic features used by ML to identify six distinct nuclei classes. Radar plots show z-scored relative levels of morphological features (nuclear area, perimeter, solidity and form factor) and DNA staining intensity (total DAPI signal). **H.** Example images of nuclei from the six classes identified by ML. Blue color shows DNA staining intensity and red color 5-ethynyl-2’-deoxyuridine (EdU) staining intensity to identify cells undergoing replication. Represented nuclei are enlarged for visualization and do not reflect actual sizes **I.** Principal component analysis (PCA) of five interphase classes based on 1,753 protein groups after data filtering. Replicates of classes are highlighted by ellipses with a 95% confidence interval. **J.** Number of differentially expressed proteins compared to unclassified nuclei (bulk). Proteins with an FDR less than 0.05 were considered significant.

To address if morphological differences between nuclei are also reflected in their proteomes, we used an unsupervised phenotype finder model to identify groups of morphologically distinct nuclei based on nuclear area, perimeter, form factor, solidity and DNA staining intensity (Fig. 3E). ML found three main nuclei classes (27-37% each) and also discovered three rare ones (2-4% each) (Fig. 3F). The resulting six distinct nuclei classes had visible differences in size and shape. Class 1 represented mitotic states while the remaining five were in interphase with varying feature weighting (Fig. 3G, H). For subsequent analysis, we focused on those five nuclei classes of unknown state. In principal component analysis (PCA), replicates of the respective proteomes clustered closely and the more frequent classes (2, 3 and 5) grouped together (Fig. 3I). The rare classes 4 and 6, which were mainly driven by their unique morphologies (large nuclei (> 4n) and ‘bean-shaped’, respectively) separated in component 1 and 2 from this group. To verify and quantify this observation, we compared each cell class proteome to a proteome of all nuclei in a field of view. This revealed that the rarest cell classes had the largest numbers of differentially expressed proteins compared to unclassified proteomes (Fig. 3J, S2B). These results demonstrate that the differences visible by microscopy translate into quantifiable proteomic differences and highlights that subcellular phenotypes are linked to distinct proteome profiles.

We next asked if the proteomic differences across the five nuclei classes could give clues to the functional differences between the interphase states (Fig. 3E, H). The 361 significantly differentially expressed proteins across classes were enriched for nuclear and cell cycle related proteins (e.g. ‘DNA unwinding involved in replication’ and ‘condensation of prophase chromosomes’), implicating the cell cycle as a strong biological driver (Fig. 4A, B). To confirm this, we compared our data to a single-cell imaging dataset including 574 cell cycle regulated proteins ^19^ and found that they were 2.3-fold < 10^−6^). In addition, nuclear area, one of the driving features between our classes, increases from G1 to S/G2 cells during interphase (Fig. 3G, Fig. S3A-C), supporting the importance of the cell cycle in defining the nuclei classes.

**Fig.4:**
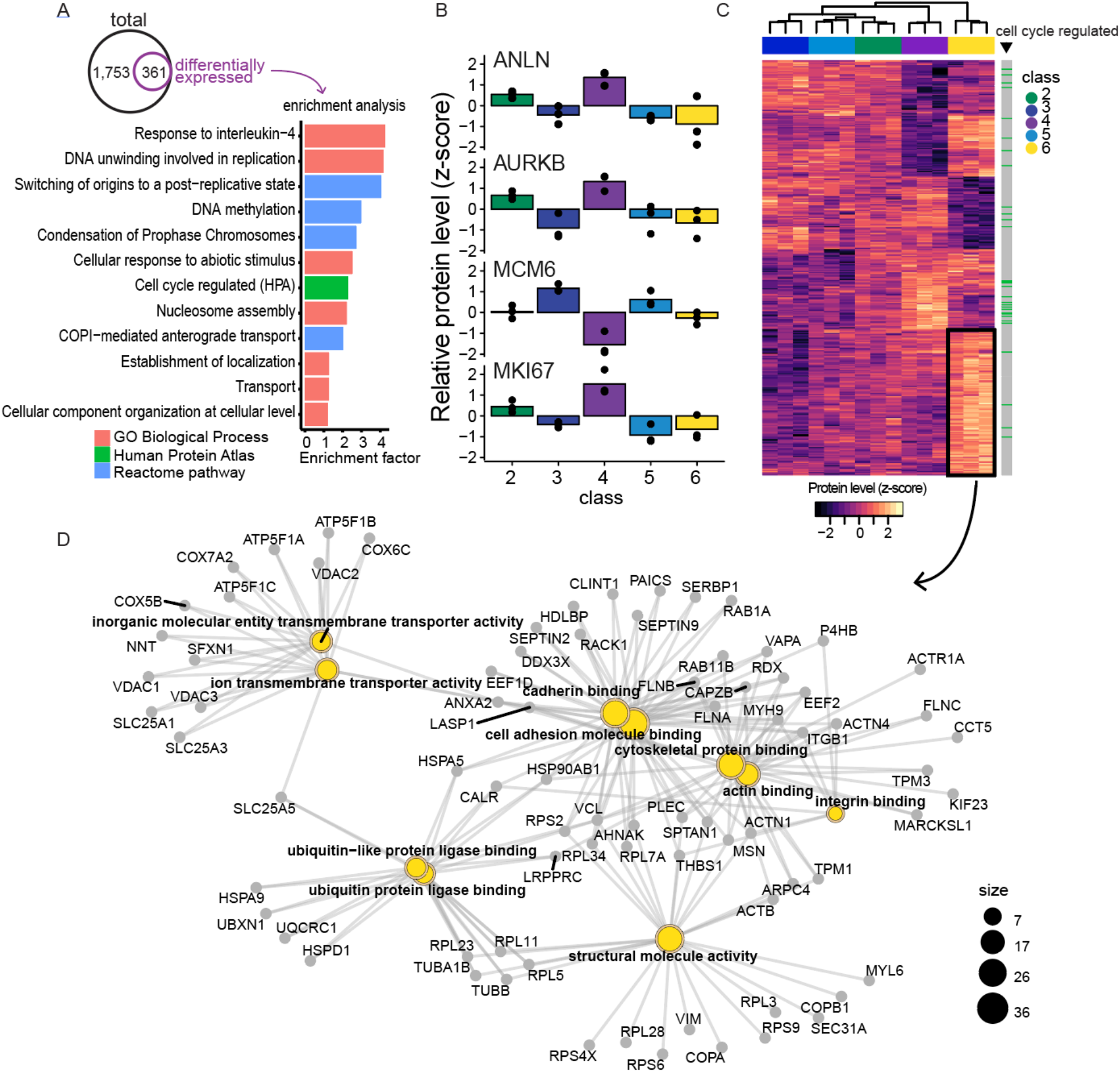
DVP links cellular phenotypes to functional protein networks in U2OS cancer cells. **A.** Bioinformatic enrichment analysis of proteins regulated between the five nuclei classes. Significant proteins (361 ANOVA significant, FDR < 0.05, *s*_*0*_ = 0.1) were compared to the set of unchanged proteins based on Gene Ontology Biological Process (GOBP), Reactome pathways and cell cycle regulated proteins reported by the Human Protein Atlas20. A Fisher’s exact test with a Benjamini-Hochberg FDR of 0.05 was used. **B.** Relative protein levels (z-score) of known cell cycle markers across the five nuclei classes. **C.** Unsupervised hierarchical clustering of all 361 ANOVA significant protein groups. Cell cycle regulated proteins reported by the HPA are shown in green in the right bar. Nuclei classes are shown in the column bar. Proteins upregulated in class 6-nuclei (yellow) are indicated by the black box. **D.** Enrichment analysis of proteins upregulated in class 6-nuclei (black box in panel C). Relationship between the top 10 most significantly enriched GO terms and proteins are shown. Node sizes represent the number of genes in each category.

We were intrigued that our single cell type proteomes contained a number of uncharacterized proteins, offering an opportunity to associate them with a potential cellular function. Focusing on three open reading frame (ORF) proteins remaining after data filtering, two of them - C7orf50 and C1orf174 - showed class specific expression patterns (p < 0.01, Fig. S3D). C7orf50 was most highly expressed in the nucleoli of classes 2 and 4 nuclei, which showed S/G2 specific characteristics (Fig. 3H, S3D, E), suggesting that its expression is cell cycle regulated. Indeed, we confirmed higher levels of C7orf50 in G1/S and S/G2 compared to G1 phase cells (Fig. S3E). As cell cycle regulated proteins tend to be over-represented in cancer prognostic associations^19^, we investigated C7orf50 in the human pathology atlas^20^ and found high expression was associated with favorable outcome in pancreatic cancer (Fig. S3G, p < 0.001). Its interaction, co-expression and co-localization with the protein LYAR (‘Cell Growth-Regulating Nucleolar Protein’) suggests a functional link between them and a potential role in cell proliferation and cancer growth (Fig. S3F, H).

Although our data revealed the cell cycle to be a strong driver of nuclei classification, class 6 showed a pronounced proteomic signature independent of known cell cycle markers (Fig. 4C, D). In these rare, bean-shaped nuclei, cytoskeletal and cell adhesion proteins (e.g. VIM, TUBB, ACTB and ITGB1) were upregulated, suggesting that they derived from migrating cells undergoing nuclear deformation, reminiscent of what has recently been described in the context of cell invasion^21, 22^. Note that we classified nuclei from 2D images but laser cutting isolates them in 3D, thus samples also probe morphology-driven protein re-localization around the nucleus as exemplified by class 6 nuclei.

These cell culture experiments establish that DVP correlates cellular phenotypes, heterogeneity and dynamics with the proteome level in an unbiased way. Common and rare phenotypes can be studied in their natural environment, preserving the native biological variation within cancer cells to link phenotypes to proteomic make-up.

## DVP applied to cancer tissue heterogeneity

We next explored if DVP could provide unbiased proteomic profiling of distinct cell classes in their individual spatial environments at high resolution. Billions of patient samples are collected routinely during diagnostic workup and biobanked in the archives of pathology departments around the world^23^. Therefore, the precise proteomic characterization of single cells in their spatial and subcellular context from these samples could have great clinical impact as an extension of the emerging field of digital pathology^24^. We selected archived tissue of a salivary gland acinic cell carcinoma, a rare and understudied malignancy of the epithelial secretory cells of the salivary gland that arises from their epithelial secretory cells. First, we developed an immunohistochemical staining protocol on glass membrane slides for routine histopathology and stained the tissue for EpCAM to outline the cellular boundaries for segmentation and feature extraction by BIAS (Methods). This revealed normal appearing and neoplastic regions with different cellular composition. While the former was mainly comprised of acinar, duct and myoepithelial cells, the carcinoma was dominated by uniform tumor cells with round nuclei and abundant basophilic cytoplasm (Fig. 5A, B).

**Fig.5:**
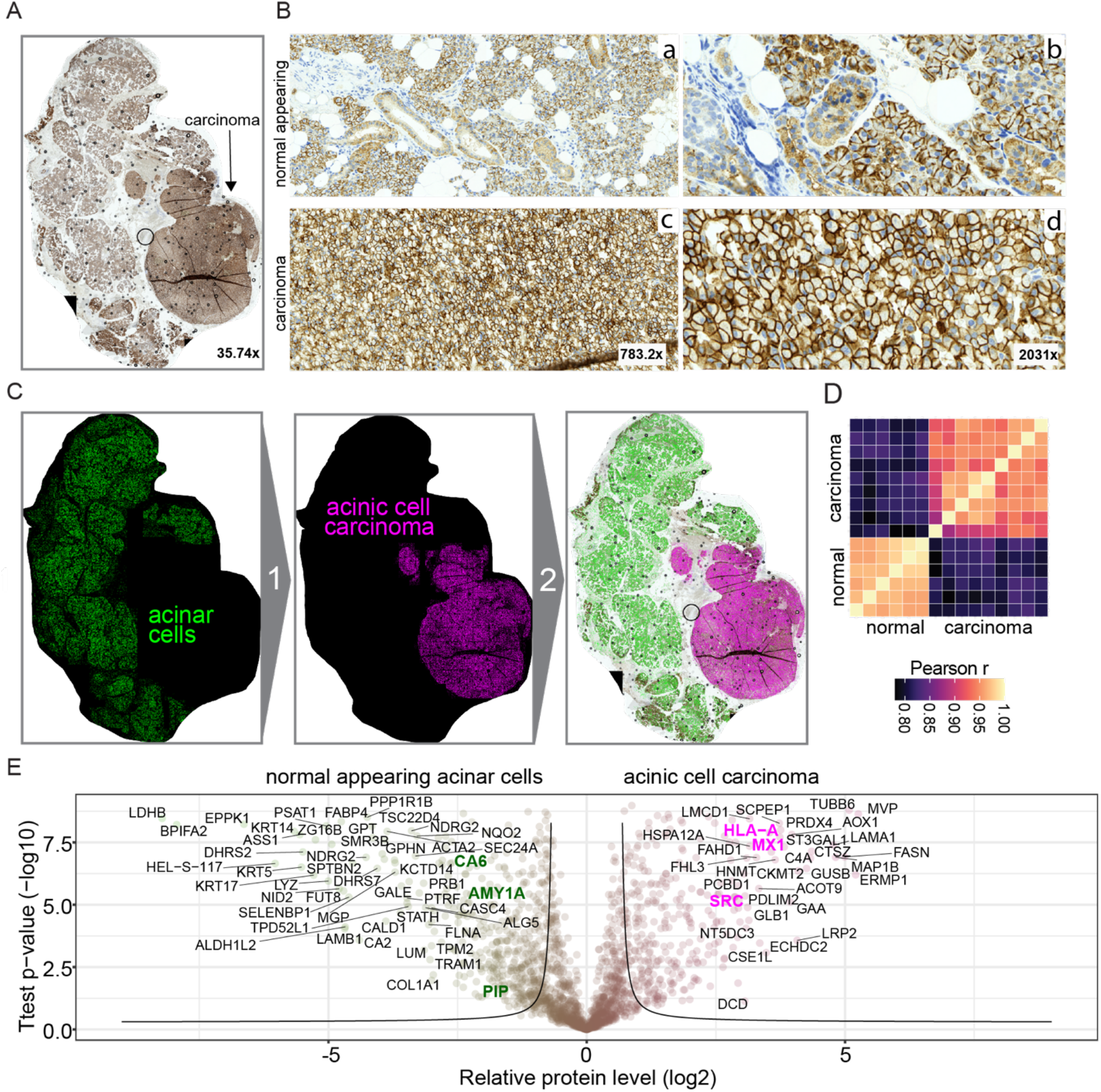
DVP applied to archived tissue of a rare salivary gland carcinoma. **A.** Immunohistochemical staining of an acinic cell carcinoma of the parotid gland by the cell adhesion protein EpCAM. **B.** Representative regions from normal appearing tissue (upper panels a and b) and acinic cell carcinoma (lower panels c and d) from A. **C.** DVP workflow applied to the acinic cell carcinoma tissue. Deep learning-based single cell detection of normal appearing (green) and neoplastic (magenta) cells positive for EpCAM. Cell classification based on phenotypic features (form factor, area, solidity, perimeter, EpCAM intensity). **D.** Proteome correlations of replicates from normal appearing (normal, n=6) or cancerous regions (cancer, n=9). **E.** Volcano plot of pairwise proteomic comparison between normal and cancer tissue. T-test significant proteins (FDR < 0.05, *s*_*0*_ = 0.1) are highlighted by black lines. Proteins higher in normal tissue are highlighted in green on the left of the volcano including known acinic cell markers (AMY1A, CA6, PIP). Proteins higher in the acinic cell carcinoma are on the right in magenta, including the proto-oncogene SRC and interferon-response proteins (MX1, HLA-A).

To identify disease specific protein signatures, we wished to directly compare the normal appearing acinar cells with the malignant cells, rather than admixing it with varying proportions of unrelated cells. To this end, we classified acinar and duct cells from normal parotid gland tissue based on their cell-type specific morphological features and isolated single cell classes for proteomic analysis (Fig. 5C and S4A). Bioinformatics of the resulting proteomes revealed strong biological differences between these neighboring cell types, reflecting their distinct physiological functions. Acinar cells, which produce and secrete saliva in secretory granules, showed high expression of proteins related to vesicle transport and glycosylation along with known acinar cell markers such as α-amylase (AMY1A), CA6 and PIP (Fig. S4B). In contrast, duct cells, which are rich in mitochondria to deal with the energy demand for transcellular saliva secretion^25^, indeed expressed high levels of mitochondria and metabolism related proteins (Fig. S4B). For comparison, we exclusively excised malignant and benign acinar cells from the various regions within the same tissue section. Interestingly, the proteomes of acinar cells clustered together regardless of disease state, demonstrating a strong cell of origin effect (Fig. S4C). Building on this foundation, we analyzed six replicates of normal appearing and nine of neoplastic regions, which showed excellent within-group quantitative reproducibility (Pearson r > 0.96). Correlation between normal and cancer were lower, reflecting disease and cell-type specific proteome changes (r = 0.8, Fig. 5D, E). Acinar cell markers in the carcinoma were significantly downregulated, consistent with previous reports^25^. We discovered an up-regulation of interferon-response proteins (e.g. MX1, HLA-A, HLA-B, SOD2) and the proto-oncogene SRC, a well-known therapeutic target^26^, together with a treasure-trove of other proteins. This highlights the ability of DVP to discover disease-specific and therapeutically relevant proteins on the basis of cell-type resolved tissue proteomics.

To investigate if DVP could resolve different states of the same cell type, we next applied it to melanoma, a highly aggressive and heterogeneous cancer associated with poor outcomes in advanced stages^27,28^. We chose an FFPE-preserved tissue specimen archived 18 years ago, as we and others have found such tissues to be amenable to MS-based proteomics, a key advantage over transcriptomics^16,29^.

Melanoma heterogeneity is driven by distinct tumor cell subpopulations and interactions with their tumor microenvironment (TME or tumor stroma), influencing disease progression, treatment response and patient survival^30^. We therefore asked if DVP could identify disease-relevant proteome signatures by comparing cancer cells from the inner tumor mass to those that were in close proximity to the stroma (Fig. 6A). To profile only melanoma cells, we isolated cells double-positive for the melanoma markers SOX10 and CD146 (Fig. 6A). Proteomes from the central tumor (central) and tumor-stroma border (peripheral) region were distinct in unsupervised hierarchical clustering and PCA (Fig. 6B). Peripheral cells showed strong BRAF, integrin and immune system related signatures, whereas in central cells up-regulated proteins with functions in DNA replication, regulation of p53 activity and mRNA splicing (FDR < 0.05, Fig. 6C, D). Prognostically relevant genes for subcutaneous melanoma have previously been reported by transcriptomics but without spatial context^31,32^. When we interrogated these markers in our data sets, we found that the outer region had significantly up-regulated favorable prognostic proteins for immune related processes (e.g. HLA-B, TAPBP, p-value < 0.01). In contrast, unfavorable proteins showed the inverse trend with higher levels in the central region (e.g. MCM3/6, CDK11A/B, DHX9) (Fig 6E, p-value < 10^−5^). Furthermore, our differential analysis highlighted ‘signaling by GCPR’, ‘extracellular matrix organization’ and a number of other relevant proteins associated with these and other cancer-relevant functions. These results show the power of DVP to quantify the spatial variability of the disease-related proteome and suggests the potential to improve molecular disease subtyping to guide clinical decision-making.

**Fig.6:**
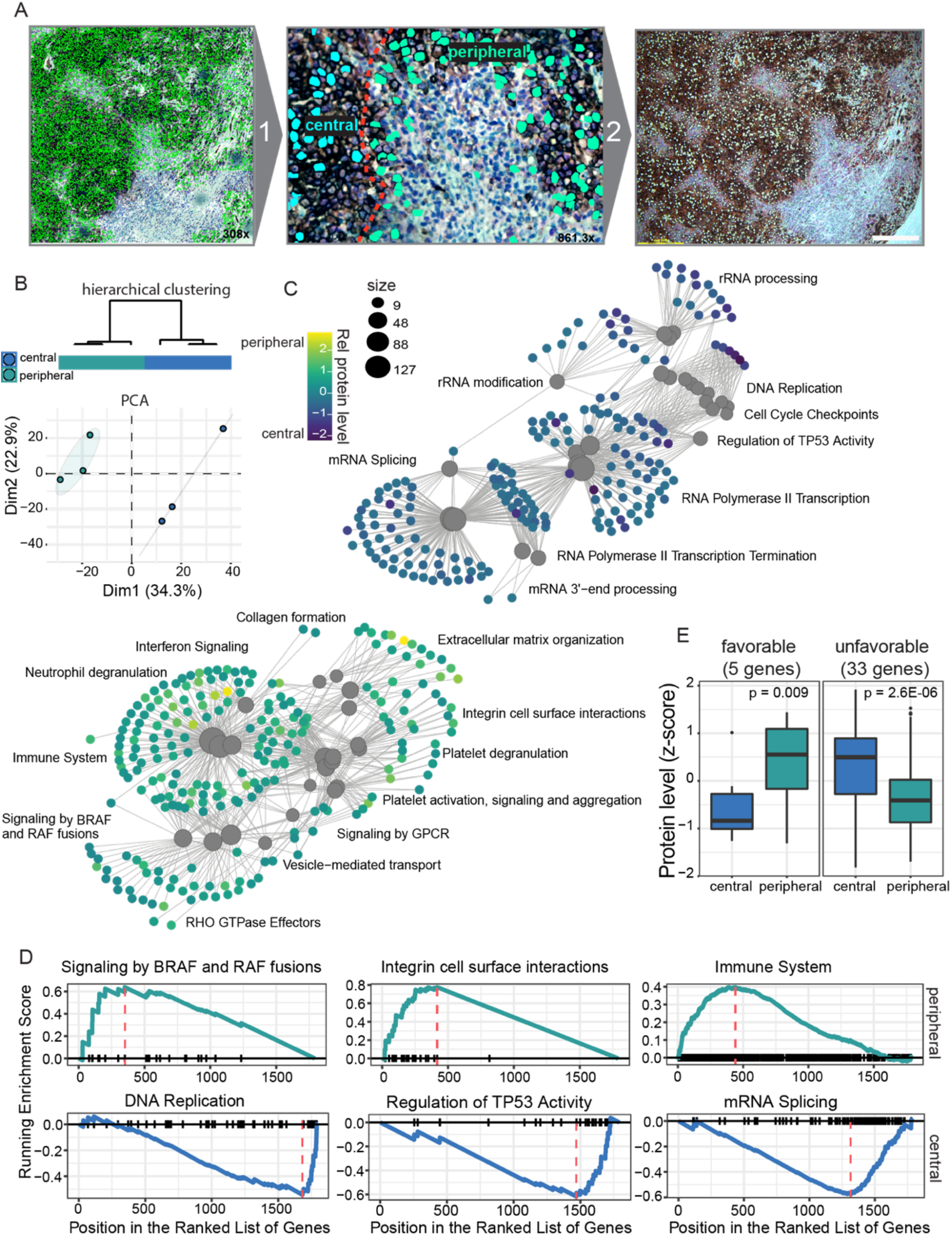
DVP applied to archived melanoma tissue. **A.** DVP applied to melanoma immunohistochemically stained for the melanoma markers SOX10 and CD146. After AI-guided single cell segmentation (left panel), SOX10 and CD146 double-positive melanoma cells were classified according to proximity to the tumor stroma (middle panel). Cells from the central tumor mass (central, marked in turquoise) and close to the tumor stroma (peripheral, marked in light green) were isolated by automated LMD (right panel) and subjected to MS-based proteomics. **B.** Upper panel: Dendrogram of unsupervised hierarchical clustering of central (n = 3) and peripheral (n = 3) melanoma cells. Lower panel: Principal component of central and peripheral melanoma cell proteomes. **C.** Enrichment analysis of proteins differentially regulated between central and peripheral melanoma cells. Relationship between all significantly enriched GO terms and proteins are shown. Node sizes represent the number of genes in each category. Node colors indicate relative log2 transformed protein levels. **D.** Gene Set Enrichment Analysis (GSEA) plot of significantly enriched pathways for central and peripheral cells. **E.** Relative protein levels comparing central and peripheral cells for favorable (left, p-value < 0.009) and unfavorable (right, p-value < 10^−5^) genes reported by^31,32^.

## Outlook

The DVP pipeline combines high resolution microscopy with new developments for image recognition, automated laser microdissection and ultra-sensitive MS-based proteomics in a robust way. Our examples demonstrate a wide range of applications, from cell culture to pathology and in principle, any biological systems that can be microscopically imaged is amenable to DVP.

A single slide can encompass hundreds of thousands of cells or more and many such slides could rapidly be scanned to isolate very rare cell states or interactions between cells. Likewise, DVP should be uniquely suited to study the proteomic composition and post-translational modifications in the extracellular matrix surrounding particular cell constellations. The resolution of excision is limited to the width of a laser beam, which is sufficient to excise individual chromosomes^33^, but cells could also be interrogated by super-resolution microscopy or highly-multiplexed imaging to better delineate precise and subtle cell states as part of their classification.

In conclusion, DVP marries increasingly powerful imagining technologies with unbiased proteomics, with a plethora of applications in basic biology and biomedicine. At the conceptual level, this technology integrates cell biology as studied by microscopy with unbiased ‘omics’ type analyses and in particular MS-based proteomics. Furthermore, the visual information gives specific context that is helpful in interpreting the proteomics data. For the field of oncology, DVP encompasses digital pathology but integrates and extends the information in stainings against a few pre-defined markers to thousands of proteins making up a cellular context.

## ACKNOWLEDGMENT

The authors thank Martin Rykær, Jeppe Madsen (NNF CPR Mass Spectrometry Platform, University of Copenhagen) and Lylia Drici (NNF CPR Proteomics Program) as well as Johannes Mueller (MPIB Munich) for technical assistance. We acknowledge Florian Hoffmann, Christoph Greb, and Falk Schlaudraff from Leica for technical support and Ede Migh, Tivadar Danka, and Mária Kovács for fruitful scientific discussions. Thomas Hartig Braunstein, Pablo Hernandez-Varas, and Clara Prats from the Core Facility of Integrated Microscopy for microscopy support. We thank Jiri Lukas for scientific support and guidance.

This work was supported by grants from the Novo Nordisk Foundation (grant agreement NNF14CC0001 and NNF15CC0001) and The Max-Planck Society for the Advancement of Science, and the Chan-Zuckerberg Initiative for partial funding of the cell cycle work (grant CZF2019-002448) to E.L., M.M. and P.H. F.C acknowledges the European Union’s Horizon 2020 research and innovation program (Marie Skłodowska-Curie individual fellowship under grant agreement (846795). BDA acknowledges support from the Lundbeck Foundation (R252-2017-1414) and the Novo Nordisk Foundation (NNF20OC0065720). PH, RH, FK and AK acknowledge support from the LENDULET-BIOMAG Grant (2018-342), from the European Regional Development Funds (GINOP-2.3.2-15-2016-00006, GINOP-2.3.2-15-2016-00026, GINOP-2.3.2-15-2016-00037), from the H2020 (ERAPERMED-COMPASS, DiscovAIR).

We acknowledge Dr. Sayuri Ito and Dr. Hisao Masai (Tokyo Metropolitan Institute of Medical Science) for providing the stable U-2 OS FUCCI cell line. LCK-GFP plasmid was a gift from Steven Green (Addgene plasmid #61099).

## AUTHOR CONTRIBUTIONS

Conceptualization, A.M. F.C., P.H. and M.M.; Methodology, A.M., F.C., A.D.B, M.B., B.D.A, M.M.; Software, R.H., F.K., A. K. and P.H.; Investigation, A.M., F.C., R.H.,; Formal Analysis, A.M., F.C., and R.H.; Writing - Original Draft, A.M., F.C., P.H. and M.M.; Writing - Review & Editing, all authors; Resources, all authors.; Data Curation, L.M.R.G., M.B., S.N., A.M., F.C., R.H., F.K., A.K., P.H.; Visualization, A.M., F.C., and R.H.; Project Administration, A.M., and P.H.; Supervision, M.M.; Funding Acquisition, F.C., P.H., E.L., and M.M.

## COMPETING INTERESTS

P.H. is the founder and a shareholder of Single-cell technologies Ltd., a biodata analysis company that owns and develops the BIAS software.

## SUPPLEMENTARY FIGURES

**Fig. S1:**
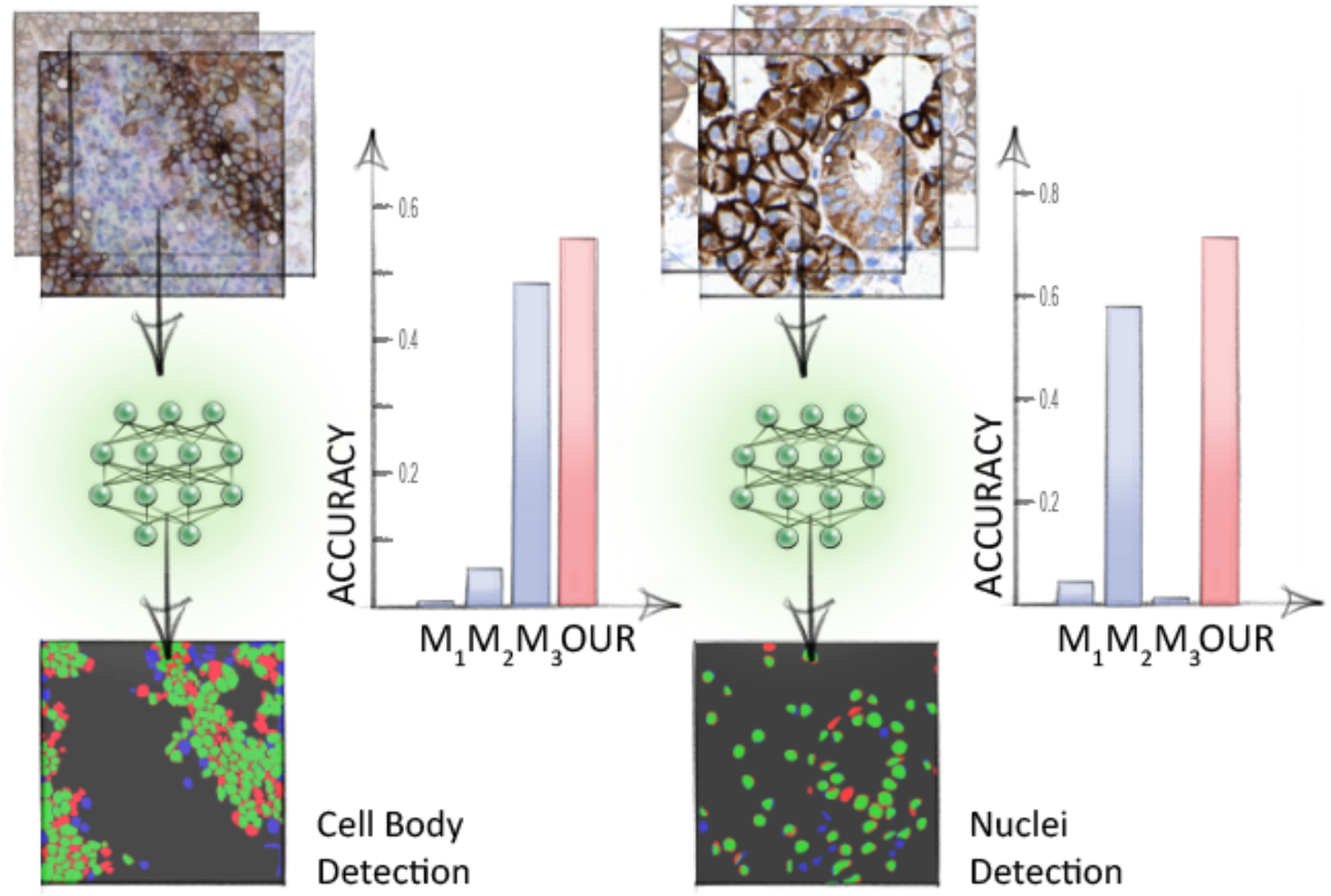
Benchmarking of segmentation algorithm. Cell body and nuclei segmentation of melanoma (left) and salivary gland tissue (right) using the Biological Image Analysis Software (BIAS). We benchmarked the accuracy of our segmentation approach using the F1 metric and compared results to three additional methods M1-M3. unet4nuclei (M_1_)^11^, conventional adaptive threshold- and object splitting-based application (M2)^13^, CellPose (M3)^12^. Visual representation of the segmentation results: green areas correspond to true positive, blue to false positive and red to false negative. Data provided in Table S1.

**Fig. S2:**
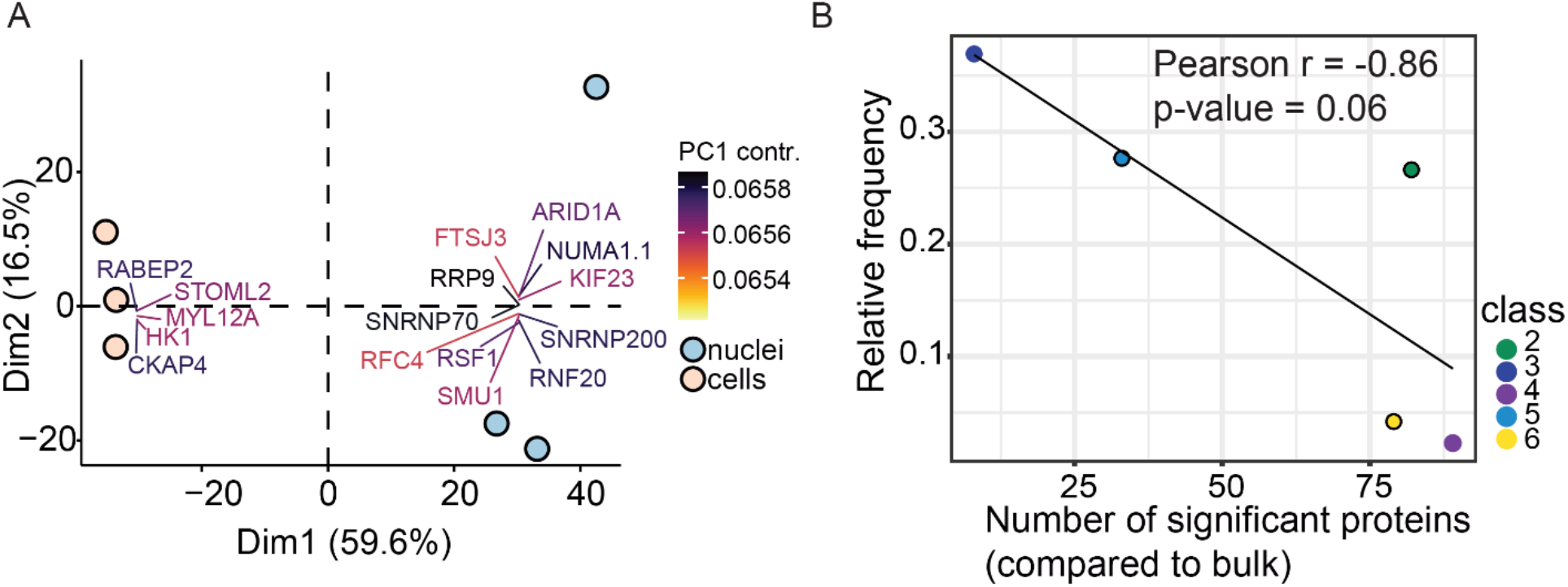
PCA and loadings of cell culture classes at sub-cellular level and number of significantly changed proteins vs. class abundance. **A.** Principal component analysis (PCA) of whole cell (n = 3) and nuclei proteomes (n = 3) based on 1,993 quantified protein groups after data filtering for no missing values. Proteins with the strongest contribution to PC1 are highlighted. **B.** Correlation between number of significantly regulated proteins per nuclei class vs relative class proportion. A linear model was fitted to the data showing an inverse correlation with Pearson r = −0.86 (p-value = 0.06).

**Fig. S3:**
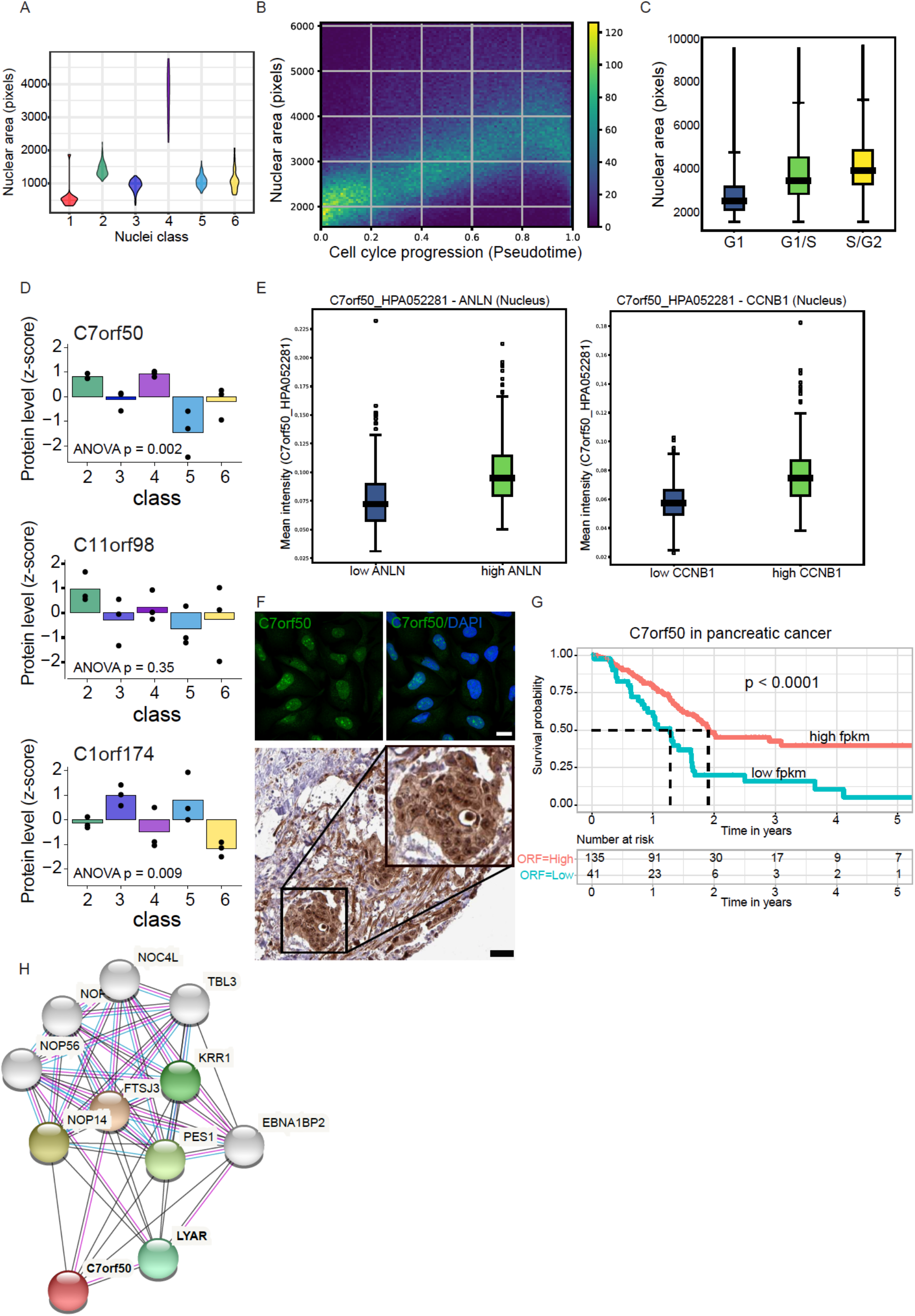
DVP discovers uncharacterized proteins with potential clinical relevance. **A.** Violin plots showing nuclear area in pixels of the 6 nuclei classes identified by ML. **B.** Nuclear area in pixels of U2OS FUCCI cells in relation to the cell cycle pseudotime19. Color code indicates point density. **C.** Nuclear area of three major cell cycle states G1, G1/S and S/G2 determined by fluorescently tagged CDT1 and GMNN intensities and Gaussian clustering. **D.** Relative protein levels of the three ORF proteins C7orf50, C11orf98 and C1orf174 across the 5 nuclei classes. C7orf50 and C1orf174 were significantly differentially regulated (p < 0.01). **E.** Mean intensities of immunofluorescently stained C7orf50 and the cell cycle markers ANLN and CCNB1 in U20S cells. C7orf50 levels were quantified in nuclei with low and high ANLN and CNNB1 intensities. **F.** Upper panel: Representative immunofluorescence images of C7orf50 and DNA (DAPI) stained U2OS cells^20,32^. Scale bar is 20 μm. Note, C7orf50 is enriched in nucleoli. Lower panel: Immunohistochemistry of a C7orf50 stained pancreatic adenocarcinoma (https://www.proteinatlas.org/ENSG00000146540-C7orf50/pathology/pancreatic+cancer#img). Image credit: Human Protein Atlas. Scale bar is 40μm. **G.** Kaplan-Meier survival analysis of pancreatic adenocarcinoma (https://www.proteinatlas.org/ENSG00000146540-C7orf50/pathology/pancreatic+cancer) based on relative C7orf50 RNA levels (FPKM, number of Fragments Per Kilobase of exon per Million reads)32. RNA-seq data is reported as median FPKM, generated by The Cancer Genome Atlas (https://www.cancer.gov/about-nci/organization/ccg/research/structural-genomics/tcga). Patients were divided into two groups based on C7orf50 levels with n=41 low and n=135 high patients. A log-rank test was calculated with p = 0.0001. **H.** String interactome analysis for C7orf50. A high confidence score of 0.7 was used with the five closest interactors highlighted by color^34^.

**Fig. S4:**
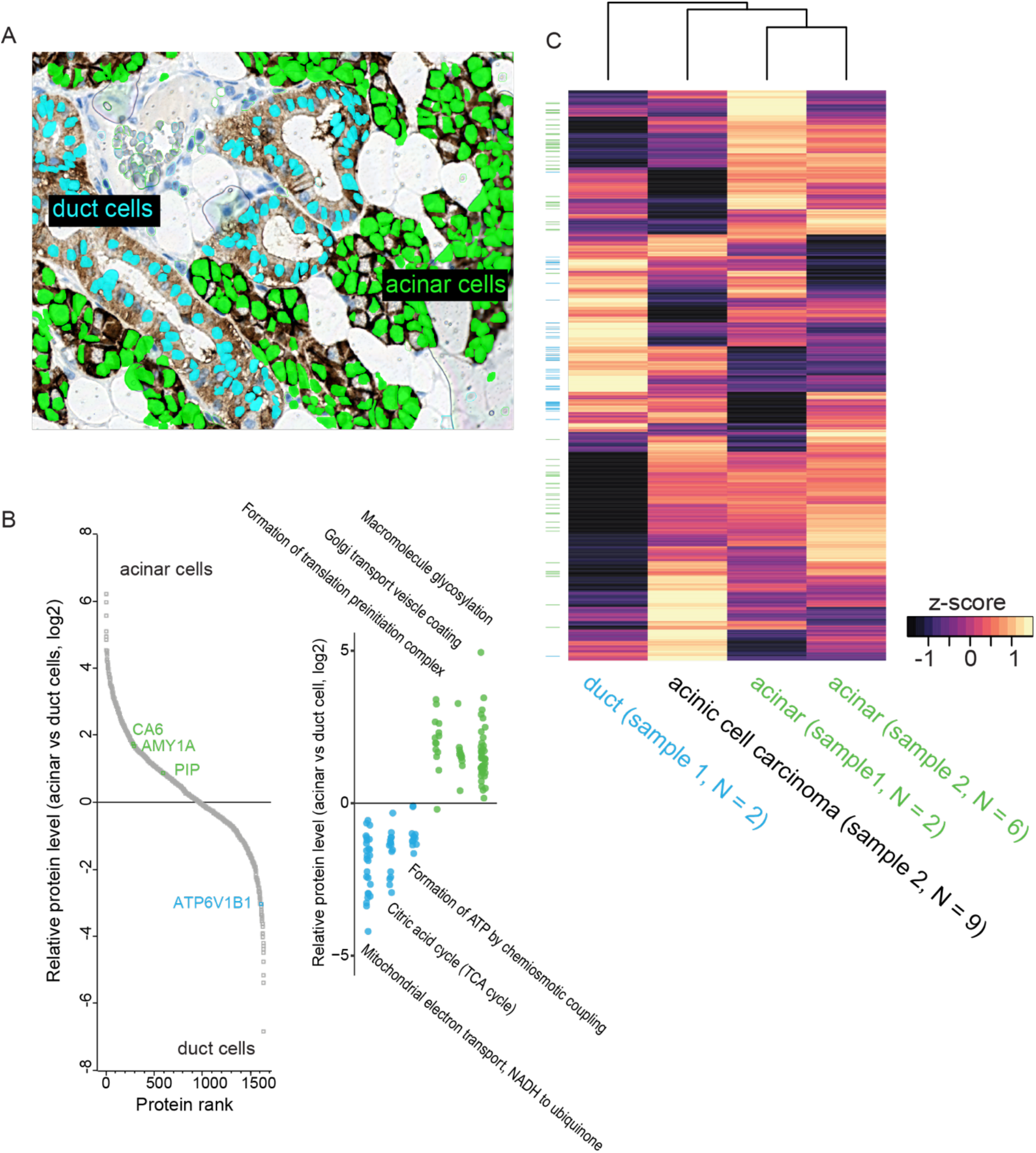
DVP applied to archival tissue of a rare salivary gland carcinoma. **A.** Immunohistochemical staining of normal salivary gland stained for the cell adhesion protein EpCAM. Supervised (random forest) ML was trained to identify acinar (green) and duct cells (turquoise). **B.** Left panel: Quantitative proteomic comparison between acinar and duct cells from tissue in A with known cell type specific markers highlighted (https://www.proteinatlas.org/humanproteome/tissue/salivary+gland). Right panel: Relative protein levels of selected pathways that were significantly higher in acinar or duct cells. **C.** Unsupervised hierarchical clustering of acinar and duct cell proteomes from two different patients together with acinar cell carcinoma cells. Note that normal acinar cells of two different tissues clustered together. Duct cells clustered furthest away. Prior to clustering, protein levels from different sample groups (duct cell tissue #1, acinar cell tissue #1, acinar cell tissue #2, carcinoma tissue #2) were averaged and z-scored. Bar on the left shows differentially expressed pathways from panel B with acini and duct specific proteins in green and turquoise, respectively.

## MATERIAL & METHODS

### Patient samples and ethics

We collected archival FFPE tissue samples of salivary gland acinic cell carcinoma and melanoma from the Department of Pathology, Zealand University Hospital, Roskilde, Denmark. The study was carried out in accordance with the institutional guidelines under approval by the local Medical Ethics Review Committee (SJ-742), the Data Protection Agency (REG-066-2019) and in agreement with Danish law (Medical Research Involving Human Subjects Act).

In accordance with the Medical Ethics Review Committee approval, all FFPE human patient tissue samples were exempted from consent as these studies used existing archived pathological specimens and not human subjects directly. Human tissue specimens were assessed by a board-certified pathologist.

### Cell lines

The human osteosarcoma cell line U2OS was grown in Dulbecco’s modified Eagle’s medium (high glucose, GlutaMAX) containing 10% FBS and penicillin-streptomycin (Thermo Fisher Scientific).

The U2OS FUCCI (Fluorescent Ubiquitination-based Cell Cycle Indicator) cells were kindly provided by Dr. Miyawaki^1^. These cells are endogenously tagged with two fluorescent proteins fused to the cell cycle regulators CDT1 (mKO2-hCdt1+) and Geminin (mAG-hGem+). CDT1 accumulates during the G1 phase while Geminin accumulates in S and G2 phases allowing cell cycle monitoring. The cells were cultivated at 37 °C in a 5.0 % CO2 humidified environment in McCoy’s 5A (modified) medium GlutaMAX supplement, (Thermo Fisher, 36600021, MA, USA) supplemented with 10% fetal bovine serum (FBS, VWR, Radnor, PA, USA) without antibiotics.

U2OS cells stably expressing a membrane-targeted form of eGFP were generated by transfection with plasmid Lck-GFP (Addgene #61099^2^) and culturing in selection medium (DMEM medium containing 10% FBS, Penicillin-Streptomycin, 400 μg/ml Geneticin) under conditions of limited dilution to yield single colonies. A clonal cell line with homogenous and moderate expression levels of Lck-eGFP at the plasma membrane was established from a single colony.

All cell lines were tested for mycoplasma (MycoAlert, Lonza) and authenticated by STR profiling (IdentiCell Molecular Diagnostics).

### Tissue preparation for immunohistochemistry

Immunohistochemical staining on membrane slides:

Membrane PEN slides 1.0 (Zeiss, Göttingen, Germany, cat. #415190-9041-000) were treated with UV light for 1 h and coated with APES (3-aminopropyltriethoxysilane) using Vectabond reagent (Vector Laboratories, Burlingame, CA, USA, cat. #SP-1800-7) according to the manufacturer’s protocol. FFPE tissue sections were cut (2.5 μm), air-dried at 37 °C overnight and heated at 60 °C for 20 min to facilitate better tissue adhesion. Next, sections were deparaffinized, rehydratrated and loaded wet on the fully automated instrument Omnis (Dako, Glostrup, Denmark). Antigen retrieval was conducted using Target Retrieval Solution pH 9 (Dako, cat # S2367) diluted 1:10 and heated for 60 minutes at 90 °C. Single stain for EpCAM (Nordic Biosite, Copenhagen K, Denmark, clone BS14, cat. #BSH-7402-1, dilution 1:400) and sequential double stain for SOX10/CD146 (SOX10; Nordic Biosite, clone BS7 cat. #BSH-7959-1, dilution 1:200 and CD146; Cell Marque, Rocklin, CA, USA, clone EP54, cat. #AC-0052, dilution 1:400) was performed and slides were incubated for 30 min (32 °C). After washing and blocking of endogenous peroxidase activity, the reactions were detected and visualized using Envision FLEX+ High pH kit (Dako, cat #GV800+GV809/GV821) according to manufacturer’s instructions. In the double stain, Envision DAB+ (Dako, cat # GV825) and Envision Magenta (Dako, cat. #GV900) Substrate Chromogen Systems was used for visualization of CD146 and SOX10, respectively. Finally, slides were rinsed in water, counterstained with Mayer’s hematoxylin, and air-dried without mounting.

### Immunofluorescence staining

Cells were first incubated with 5-ethynyl-2 deoxyuridine (EdU) for 20 min, then fixed for 5 min at room temperature with 4 % paraformaldehyde and washed three times with PBS. Cells were then permeabilized with PBS/0.2% Triton-X for 2 min on ice and washed three times with PBS. Cells were then stained with an EdU labeling kit (Life Technologies) and counterstained with Hoechst 33342 for 10 min. Slides were mounted with GB mount (GBI Labs # E01-18).

96-well glass bottom plates (Greiner Sensoplate Plus, Greiner Bio-One, Germany) were coated with 12.5 μg/ml human fibronectin (Sigma Aldrich, Darmstadt, Germany) for 1 h at RT. Immunocytochemistry was carried out following an established protocol^3^. 8,000 U2OS cells were seeded in each well and incubated in a 37 °C and 5% CO2 environment for 24 hours. Cells were washed with PBS, fixed with 40 μl 4% ice-cold PFA and permeabilized with 40 μl 0.1 Triton X-100 in PBS for 3×5 min. Rabbit polyclonal HPA antibodies targeting the proteins of interest were diluted in blocking buffer (PBS + 4% FBS) at 2-4 μg/ml along with marker primary antibodies (see just below) and incubated overnight at 4°C. Cells were washed with PBS for 4×10 min and incubated with secondary antibodies (goat anti-rabbit Alexa488 (A11034, Thermo Fisher), goat anti-mouse Alexa555 (A21424, Thermo Fisher), goat anti-chicken Alexa647 (A21449, Thermo Fisher)) in blocking buffer at 1.25 μg/ml for 90 min at RT. Cells were counterstained in 0.05 μg/ml DAPI for 15 min, washed with for 4×10min and mounted in PBS.

Primary antibodies used:

For C7orf50 cell cycle validation:

mouse anti ANLN at 1.25 ug/ml (amab90662, Atlas Antibodies)

Mouse anti CCNB1 at 1 ug/ml: (610220, BD Biosciences)

### High-resolution microscopy

Images of immunofluorescence-labeled cell cultures were acquired using an AxioImager Z.2 microscope (Zeiss, Germany), equipped with widefield optics, a 20×, 0.8-NA dry objective, a quadruple-band filter set for Hoechst, FITC, Cy3 and Cy5 fluorescent dyes. Widefield acquisition was performed using the Colibri7 LED light source and an AxioCam 702 mono camera with 5,9 mm/px. Z-stacks with 19 z-slices were acquired at 3 mm increments to capture the optimal focus plane. Images were obtained automatically with the Zeiss Zen 2.6 (blue edition) at non-saturating conditions (12-bit dynamic range).

IHC images from salivary gland and melanoma tissue were obtained using the automated slide scanner Zeiss Axio Scan.Z1 (Zeiss, Germany) for brightfield microscopy. Brightfield acquisition was obtained using the VIS LED light source and a CCD Hitachi HV-F202CLS camera. PEN slides were scanned with a 20×, 0.8-NA dry objective yielding a resolution of 0,22 mm/px. Z-stacks with 8 z-slices were acquired at 2 mm increments to capture the optimal focus plane. Color images were obtained automatically with the Zeiss Zen 2.6 (blue edition) at non-saturating conditions (12-bit dynamic range).

#### Widefield fluorescence microscopy for validation of cell cycle dependent C7orf50 expression

Cells were imaged on a Leica Dmi8 widefield microscope equipped with a 0.80 N/A 40x air objective and a Hamamatsu Flash 4.0 V3 camera using the LAS X software. The segmentation of each cell was performed using the Cell Profiler software^4^ using DAPI for nuclei segmentation. The mean intensity of the target protein and the cell cycle marker protein was measured in the nucleus. The cells were grouped into the G1 and G2 phases of the cell cycle by using the 0.2 and 0.8 quantile of ANLN or CCNB1 intensity levels in the nucleus and cell cycle dependent expression of C7orf50 was validated by comparing differences in expression levels between G1 and G2 cells.

### Laser microdissection

To excise cells or nuclei we used the Leica LMD7 system, which we had adapted for automated single cell automation. High cutting precision was achieved using a HC PL FLUOTAR L 63x/0.70 (tissue) or 40x/0.60 (cells cultures) CORR XT objective. We used the Leica Laser Microdissection V 8.2.3.7603 software (adapted for this project) for full automated excision and collection of more than 700 contours per hour.

Leica LMD 7 cutting accuracy (Leica R&D, patent EP1276586)

For 150x objective: 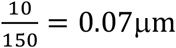

### Biological Image Analysis Software (BIAS) introduction

A typical image analysis workflow in BIAS consists of multiple images processing steps, including image preprocessing, object segmentation, contour post-processing, feature extraction and statistical analysis, supervised or unsupervised machine learning methods for phenotype classification, cell selection and cell extraction (by a selected or supported micro-dissection microscope). For each workflow step BIAS provides conveniently customizable modules.

Images were captured with a Zeiss Axio Scan.Z1 or AxioImager Z.2 microscope, both are supported by the analysis software with preservation of correct spatial information (spatial topology and size). Other types of microscopes with similar support include 3DHistech, Hamamatsu, GE IN Cell, Molecular Devices ImageXpress Micro, Leica SP, Perkin Elmer Opera and Operetta. It is also possible to import standard image files with editable image orientation and resolution. Illumination-correction algorithm, primarily CIDRE^5^ (within BIAS) were applied where it was necessary to solve the frequent ‘vignetting’ effect observable in raw microscopy images.

Image preprocessing was followed by deep learning-based nucleus and cell segmentation modules (see segmentation methods and accuracy evaluation) further refined by unary and binary morphological operators (e.g.: dilation, erosion, cavity filling, addition and subtraction). For example, subtraction can be used to calculate the cytoplasm-only region from cell and nuclei masks.

Results of different segmentation algorithms may be connected by a linking module to form complex structures, e.g. an abstract cell object might be constructed from a segmented nucleus, cytoplasm and proteins where each component can be analyzed individually or as a whole. Objects were forwarded to the feature extraction modules, configurable to extract properties from the selected image channels and cell components. A multitude of features can be retrieved from the image and contour data, such as shape (e.g.: area, perimeter, form factor, solidity etc.), intensity (e.g.: min, max, mean, total) or texture (e.g.: Haralick features) and represented in a feature matrix^6^. Features to be extracted may vary by experiments according to their specific requirements each, potentially containing up to hundreds or a few thousands per cell depending on the configuration. Features from the neighboring regions of each cell can be incorporated as well, to further improve accuracy where local neighborhoods might also contain valuable information for the cell phenotypes (such as in tissues)^7^. The resulting feature matrices can be analyzed internally or exported to a 3rd party tool^8^. Subsequently reimport of extended feature matrices into BIAS is also possible to extend the statistical capabilities or simply to visualize the data in the plate overview.

Internal analysis tools include simple, value-based statistics, manual gating, automatic feature space clustering and interactive supervised machine learning; additionally, these may also be combined. Final cell selection can depend on simple, value-based statistics or complex queries searching in multiple feature and classification matrices.

With manual gating, two features can be represented in a two-dimensional coordinate system and with cluster centers defined manually, samples are displayed at their actual position in the coordinate system. K-Means clustering can automatically find a fixed number of cluster centers in the feature space with an arbitrary number of dimensions.

During supervised machine learning, the biologist defines the phenotypes of interest and provides training samples (usually around a hundred samples for each class). Training is iterative and interactive, refinable and new phenotypes may be identified using an active learning technique^9^. A cross validation tool is provided to continuously monitor accuracy, so that when a satisfactory threshold is reached, all other cells in the whole experiment are classified. Different machine learning approaches are implemented in the BIAS software for various experimental needs. Such methods are 1) Multilayer perceptron (a feedforward artificial neural network), 2) Support-vector machine (separates the feature space by hyperplanes between the training samples), 3) Random forest (a number of decision trees trained to separate the training data into classes) and 4) Logistic regression (a statistical model that determines the probability of passing or failing the criteria of a certain class). These classification algorithms can analyze and classify tens of thousands of cells in a matter of seconds.

Results of the feature extraction and classification phases can be summarized in the statistics module, in addition it also provides an interface where custom queries can be written in SQL language and executed on the cell information database containing feature and classification data for all cells in the experiment. Cross queries between different classification and feature matrices are also supported. Query templates and wizards are provided for the most common questions.

The queries might be as simple as e.g. listing N items that have the highest value in a selected feature, or rather complex e.g. to calculate the sum and ratio of the areas of cells belonging to different classes). Results can be represented in graphs, heatmaps or used to filter cells for capturing by a suitable microscope.

The visualization tool supports a virtually unlimited number of channels with adjustable intensity window, gamma and look-up-table settings. An interactive, zoomable overview of the whole experiment (let it be a slide or a plate) can be displayed, reflecting the changes in visualization or data processing real-time. The results of all processing steps could be overlaid on it, such as segmentation masks, feature heatmaps or phenotype classification as color-coded segments, etc. A schematic display of the multi-well plate helps the navigation, enabling manual selection of cross-field areas for isolation.

The isolation and collection module uses a registration algorithm based on a marker or point-of-interest (POI) to connect the coordinate systems of the source and the isolation microscopes as well as to transfer the contour points to that of the isolation microscope. The tool supports sorting cells into different collectors and also cutting components in order (e.g. cutting nuclei into a microplate well or collection cap first then remaining cytoplasm into another). Cells of interest can be selected manually or using the statistics module. To preserve object integrity, it allows the user to define cutting offsets or exclude touching regions thereby preventing undesirable laser-induced damage.

### Segmentation methods and accuracy evaluation

NucleAIzer^10^ models were integrated into BIAS and customized for these experiments by retraining and refining the nucleus and cytoplasm segmentation models. Firstly, style transfer^11^ learning was performed as follows. Given a new experimental scenario such as our melanoma or salivary gland tissue sections stained immunohistochemically, the acquisition of which produces such an image type no annotated training data exists for, preventing efficient segmentation with even powerful deep learning methods. With an initial segmentation or manual contouring by experts (referred to as annotation) a small mask dataset is acquired (masks represent e.g. nuclei) which is used to generate new mask images such that the spatial distribution, density and morphological properties of the generated objects (e.g. nuclei) are similar to those of the annotated images. The initial masks and their corresponding microscopy images are used to train a style transfer model that learns how to generate the texture of the microscopy images on the masks marking objects: foreground to mimic e.g. nuclei and background for surrounding e.g. tissue structures. Parallelly, artificial masks of either nucleus or cytoplasm objects were created and input to the style transfer network that generated realistic-looking synthetic microscopy images with the visual appearance of the original experiment. Hence, with this artificially created training data (synthetic microscopy images and their corresponding, also synthetic masks) their applied deep learning segmentation model, Mask R-CNN, is prepared for the new image type and can accurately segment the target compartments.

We benchmarked the accuracy of the segmentation approach on a fluorescent LCK-U2OS cell line as well as tissue samples of melanoma and salivary gland, and compared results to three additional methods, including two deep learning approaches: unet4nuclei (denoted as M_1_ on Fig. 2A and S1)^12^ and CellPose (M3)^13^, alongside a widely-used, conventional adaptive threshold- and object splitting-based application (M2)^4^. We note that M_1_ is not intended for cytoplasm segmentation (see details in^12^ and below). Segmentation accuracy according to the F1 metric is displayed as bar plots (Fig. 2A, S1), while visual representation in a color-coded manner is also provided.

unet4nuclei^12^ is optimized to segment nuclei on cell culture images, and 2) CellPose^13^ is an approach intended for either nucleus or cytoplasm segmentation on various microscopy image types, while CellProfiler^4^ is a conventional threshold- and object splitting-based software broadly used in the bioimage analysis community. unet4nuclei as its name suggests is primarily intended for nucleus segmentation and uses a U-Net-based network after pre-processing of input images, then post-processes detected objects. Cellpose uses a vector flow-representation of instances and its neural network (also based on U-Net) predicts and combines horizontal and vertical flows. unet4nuclei has successfully been applied in nucleus segmentation of cell cultures, while CellPose is able to generalize well on various image modalities even outside microscopy and can be used to segment nuclei and cytoplasms.

We evaluated our segmentation performance (and comparisons) according to the F1-score metric calculated at 0.7 IoU (intersection over union) threshold. IoU also known as Jaccard index was calculated from the overlapping region of the predicted (segmented) object with its corresponding ground truth (real) object at a given threshold (see formulation below). True positive (TP), false positive (FP) and false negative (FN) objects were counted accordingly, if they had IoU greater than the threshold *t* (in our case 0.7), to yield the F1-score at this threshold (see formulation below). Considering the mean F1-scores measured, we conclude that the applied deep learning-based segmentation method^10^ available in BIAS produced segmentations on both nucleus and cytoplasm level in a higher quality than the compared methods; see results on Fig. 2A and S1.

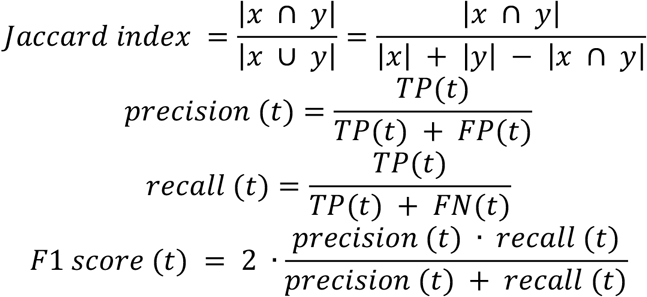

Our evaluation results of nucleus and cell body segmentation on melanoma-, salivary gland tissues and U2OS cells is presented in (Supplementary Table 1).

**Supplementary Table 1.** F1-scores of the compared segmentation methods on our samples. The methods are as follows: M1 is unet4nuclei^12^, M2 is CellProfiler^4^, M3 is Cellpose^13^, while OUR refers to nucleAIzer^10^ (implemented in BIAS). High scores are highlighted in bold. Asterisks mark that M1 is intended for nucleus segmentation but was applied to segment cytoplasm.

| sample | method |  |  |  |
| --- | --- | --- | --- | --- |
|  | M1 | M2 | M3 | OUR |
| U2OS cyto | 0.0667* | 0.5994 | 0.7205 | **0.7336** |
| Melanoma nuc | 0.1126 | 0.4386 | 0.1801 | **0.5498** |
| Melanoma cyto | 0.0058* | 0.0549 | 0.4859 | **0.5536** |
| Salivary gland nuc | 0.0476 | 0.5830 | 0.0160 | **0.7158** |
| Salivary gland cyto | 0.0991* | 0.0703 | 0.2312 | **0.4111** |

### Sample preparation for mass spectrometry

Cell culture (nuclei or whole cells) and tissue samples were collected by automated laser microdissection into 384-well plates (Eppendorf 0030129547). For the collection of different U2OS nuclei classes (Fig. 3 and 4), we normalized nuclear size differences (resulting in different total protein amounts) by the number of collected objects per class. On average, we collected 267 nuclei per sample. For FFPE tissue samples of salivary gland and melanoma, (2.5 μm thick section cut in microtome) an area of 80,000 – 160,000 μm2 per sample was collected, an estimated number of 100-200 cells based on the average HeLa cell volume of 2,000 μm3 (BNID 100434).

20μl of ammonium bicarbonate (ABC) were added to each sample well and the plate closed with sealing tape (Corning, CLS6569-100EA). Following vortexing for 10 s, plates were centrifuged for 10 min at 2000g and heated at 95C for 30 min (cell culture) or 60 min (tissue) in a thermal cycler (Biorad S1000 with 384-well reaction module) at a constant lid temperature of 110 °C. 5 μl 5x digestion buffer (60% acetonitrile in 100 mM ABC) was added and samples heated at 75 °C for another 30 min. Samples were shortly cooled down and 1 μl LysC added (pre-diluted in ultra-pure water to 4 ng/μl) and digested for 4 h at 37 °C in the thermal cycler. Subsequently, 1.5 μl trypsin was added (pre-diluted in ultra-pure water to 4ng/μl) and incubated overnight at 37 °C in the thermal cycler. Next day, digestion was stopped by adding trifluoroacetic acid (TFA, final concentration 1% v/v) and samples vacuum-dried (approx. 1.5 h at 60 °C). 4 μl MS loading buffer (3% acetonitrile in 0.2% TFA) was added, the plate vortexed for 10s and centrifuged for 5 min at 2000g. Samples were stored at −20 °C until LC-MS analysis.

### High-pH reversed-phase fractionation

We used high-pH reversed-phase fractionation to generate a deep U2OS cell precursor library for data-independent (DIA) MS analysis (below). Peptides were fractionated at pH 10 with the spider-fractionator^14^. 30 μg of purified peptides were separated on a 30 cm C18 column in 100 min and concatenated into 12 fractions with 90 s exit valve switches. Peptide fractions were vacuum-dried and reconstituted in MS loading buffer for LC-MS analysis.

### LC-MS analysis

Liquid chromatography mass spectrometry (LC-MS) analysis was performed with an EASY-nLC-1200 system (Thermo Fisher Scientific) connected to a modified trapped ion mobility spectrometry quadrupole time-of-flight mass spectrometer with about five-fold higher ion current^15^ (timsTOF Pro, Bruker Daltonik GmbH, Germany) with a nano-electrospray ion source (Captive spray, Bruker Daltonik GmbH). The autosampler was configured for sample pick-up from 384-well plates.

Peptides were loaded on a 50 cm in-house packed HPLC-column (75μm inner diameter packed with 1.9μm ReproSilPur C18-AQ silica beads, Dr. Maisch GmbH, Germany).

Peptides were separated using a linear gradient from 5-30% buffer B (0.1% formic acid, 80% ACN in LC-MS grade H2O) in 55 min followed by an increase to 60% for 5 min and 10 min wash at 95% buffer B at 300nl/min. Buffer A consisted of 0.1% formic acid in LC-MS grade H2O. The total gradient length was 70 min. We used an in-house made column oven to keep the column temperature constant at 60 °C.

Mass spectrometric analysis was performed essentially as described in Brunner et al.^15^, either in data-dependent (ddaPASEF) (Fig. 5 and 6) or data-independent (diaPASEF) mode (Fig. 3 and 4). For ddaPASEF, 1 MS1 survey TIMS-MS and 10 PASEF MS/MS scans were acquired per acquisition cycle. Ion accumulation and ramp time in the dual TIMS analyzer was set to 100 ms each and we analyzed the ion mobility range from 1/K0 = 1.6 Vs cm-2 to 0.6 Vs cm-2. Precursor ions for MS/MS analysis were isolated with a 2 Th window for m/z < 700 and 3 Th for m/z >700 in a total m/z range of 100-1.700 by synchronizing quadrupole switching events with the precursor elution profile from the TIMS device. The collision energy was lowered linearly as a function of increasing mobility starting from 59 eV at 1/K0 = 1.6 VS cm-2 to 20 eV at 1/K0 = 0.6 Vs cm-2. Singly charged precursor ions were excluded with a polygon filter (otof control, Bruker Daltonik GmbH). Precursors for MS/MS were picked at an intensity threshold of 1.000 arbitrary units (a.u.) and resequenced until reaching a ‘target value’ of 20.000 a.u taking into account a dynamic exclusion of 40 s elution. For DIA analysis, we made use of the correlation of Ion Mobility (IM) with m/z and synchronized the elution of precursors from each IM scan with the quadrupole isolation window. The collision energy was ramped linearly as a function of the IM from 59 eV at 1/K0 = 1.6 Vs cm−2 to 20 eV at 1/K0 = 0.6 Vs cm−2. We used the ddaPASEF method for library generation^16^.

### Data analysis of proteomic raw files

Mass spectrometric raw files acquired in ddaPASEF mode (Fig. 5 and 6) were analyzed with MaxQuant (version 1.6.7.0)^17,18^. The Uniprot database (2019 release, UP000005640_9606) was searched with a peptide spectral match (PSM) and protein level FDR of 1%. A minimum of seven amino acids was required including N-terminal acetylation and methionine oxidation as variable modifications. Due to omitted reduction and alkylation, cysteine carbamidomethylation was removed from fixed modifications. Enzyme specificity was set to trypsin with a maximum of two allowed missed cleavages. First and main search mass tolerance was set to 70 ppm and 20 ppm, respectively. Peptide identifications by MS/MS were transferred by matching four-dimensional isotope patterns between the runs (MBR) with a 0.7-min retention-time match window and a 0.05 1/K0 ion mobility window. Label-free quantification was performed with the MaxLFQ algorithm^19^ and a minimum ratio count of one. For diaPASEF raw file analysis (Fig. 3 and 4), we used a hybrid library approach combining a 12-fraction high-pH reversed-phase fractionated precursor library from U2OS cells with the directDIA search of the diaPASEF raw files. The hybrid library consisted of 178,948 precursors, 127,049 peptides, 9,954 protein groups and was generated with the Spectronaut software (version 14.5.200813.47784, Biognosys AG, Schlieren, Switzerland) under default settings. Search parameters were according to default settings. Protein intensities were normalized using the ‘Local Normalization’ (Q-value complete) algorithm in Spectronaut based on a local regression model. A protein and precursor FDR of 1% was used. Decoy hits and proteins, which did not pass the Q-value threshold, were filtered out prior to data analysis.

### Bioinformatic analysis

Proteomics data analysis was performed with Perseus^20^ and within the R environment (https://www.r-project.org/). MaxQuant output tables were filtered for ‘Reverse’, ‘Only identified by site modification’, and ‘Potential contaminants’ before data analysis. Data was stringently filtered to only keep proteins with 30% or less missing values (those displayed as 0 in MaxQuant output). Missing values were imputed based on a normal distribution (width = 0.3; downshift = 1.8) prior to statistical testing. Principal component analysis was performed in R. For multi-sample (ANOVA) or pairwise proteomic comparisons (two-sided unpaired t-test), we applied a permutation-based FDR of 5% to correct for multiple hypothesis testing. An *s*_*0*_ value^21^ of 0.1 was used for the pairwise proteomic comparison in Fig. 5E. Pathway enrichment analysis was performed in Perseus (Fig. 4A, Fisher’s exact test with Benjamini-Hochberg FDR of 0.05) or ClusterProfiler^22^ (Fig. 4D and 5C, D). For all ClusterProfiler analyses, an FDR filter of 0.05 was used. Minimum category size was set to 20 and maximum size to 500.

